# CRISPR adenine and cytosine base editors with reduced RNA off-target activities

**DOI:** 10.1101/631721

**Authors:** Julian Grünewald, Ronghao Zhou, Sowmya Iyer, Caleb A. Lareau, Sara P. Garcia, Martin J. Aryee, J. Keith Joung

## Abstract

CRISPR-guided DNA base editors enable the efficient installation of targeted single-nucleotide changes. Cytosine or adenine base editors (**CBE**s or **ABE**s), which are fusions of cytidine or adenosine deaminases to CRISPR-Cas nickases, can efficiently induce DNA C-to-T or A-to-G alterations in DNA, respectively^1-4^. We recently demonstrated that both the widely used CBE BE3 (harboring a rat APOBEC1 cytidine deaminase) and the optimized ABEmax editor can induce tens of thousands of guide RNA-independent, transcriptome-wide RNA base edits in human cells with high efficiencies^5^. In addition, we showed the feasibility of creating SElective Curbing of Unwanted RNA Editing (**SECURE**)-BE3 variants that exhibit substantially reduced unwanted RNA editing activities while retaining robust and more precise on-target DNA editing^5^. Here we describe structure-guided engineering of SECURE-ABE variants that not only possess reduced off-target RNA editing with comparable on-target DNA activities but are also the smallest *Streptococcus pyogenes* Cas9 (**SpCas9**) base editors described to date. In addition, we tested CBEs composed of cytidine deaminases other than APOBEC1 and found that human APOBEC3A (**hA3A**) cytidine deaminase CBE induces substantial transcriptome-wide RNA base edits with high efficiencies. By contrast, a previously described “enhanced” A3A (**eA3A**) cytidine deaminase CBE or a human activation-induced cytidine deaminase (**hAID**) CBE induce substantially reduced or near background levels of RNA edits. In sum, our work describes broadly useful SECURE-ABE and -CBE base editors and reinforces the importance of minimizing RNA editing activities of DNA base editors for research and therapeutic applications.

## Introduction

CRISPR-guided DNA base editor technology has been rapidly adopted for use in a variety of different cell types and organisms^1^. We recently demonstrated that the BE3 CBE and ABEmax can induce transcriptome-wide RNA off-target edits in human cells^5^. We also showed the feasibility of engineering a SECURE version of BE3^5^. Based on these studies, we were interested in exploring whether it might be possible to engineer a SECURE ABE and to define the RNA off-target profiles of other CBEs bearing cytidine deaminases other than the APOBEC1 enzyme present in BE3.

## Results

To attempt to engineer SECURE-ABE variants, we first used a protein truncation strategy to reduce the RNA recognition capability of the widely used ABEmax fusion. ABEmax harbors a single-chain heterodimer of the wild type (WT) *E. coli* TadA adenosine deaminase monomer (which deaminates adenines on tRNA) fused to an engineered *E. coli* TadA monomer that was modified by directed evolution to deaminate DNA adenines^3,6,7^ (**Fig. 1a**). We hypothesized that the WT TadA monomer should still be capable of recognizing its tRNA substrate and therefore might recruit ABEmax to deaminate RNA adenines that lie in the same or a similar sequence motif to that present in the tRNA. Consistent with this idea, a re-analysis of our previously published RNA-seq data^5^ revealed that adenines that are edited at the highest efficiencies (80-100%) are embedded in a more extended CU**<u>A</u>**CGAA motif, which contrasts to the shorter UA sequence observed across all edits (**Fig. 1b**). Importantly, the CUACGAA motif is a perfect match to the sequence surrounding the adenine deaminated in the tRNA substrate of the WT *E. coli* TadA enzyme^6^. We reasoned that removing the WT TadA domain from ABEmax might reduce its RNA editing activity and we therefore generated a smaller ABEmax variant lacking this domain that we refer to as **miniABEmax** (**Fig. 1a**).

**Figure 1.**
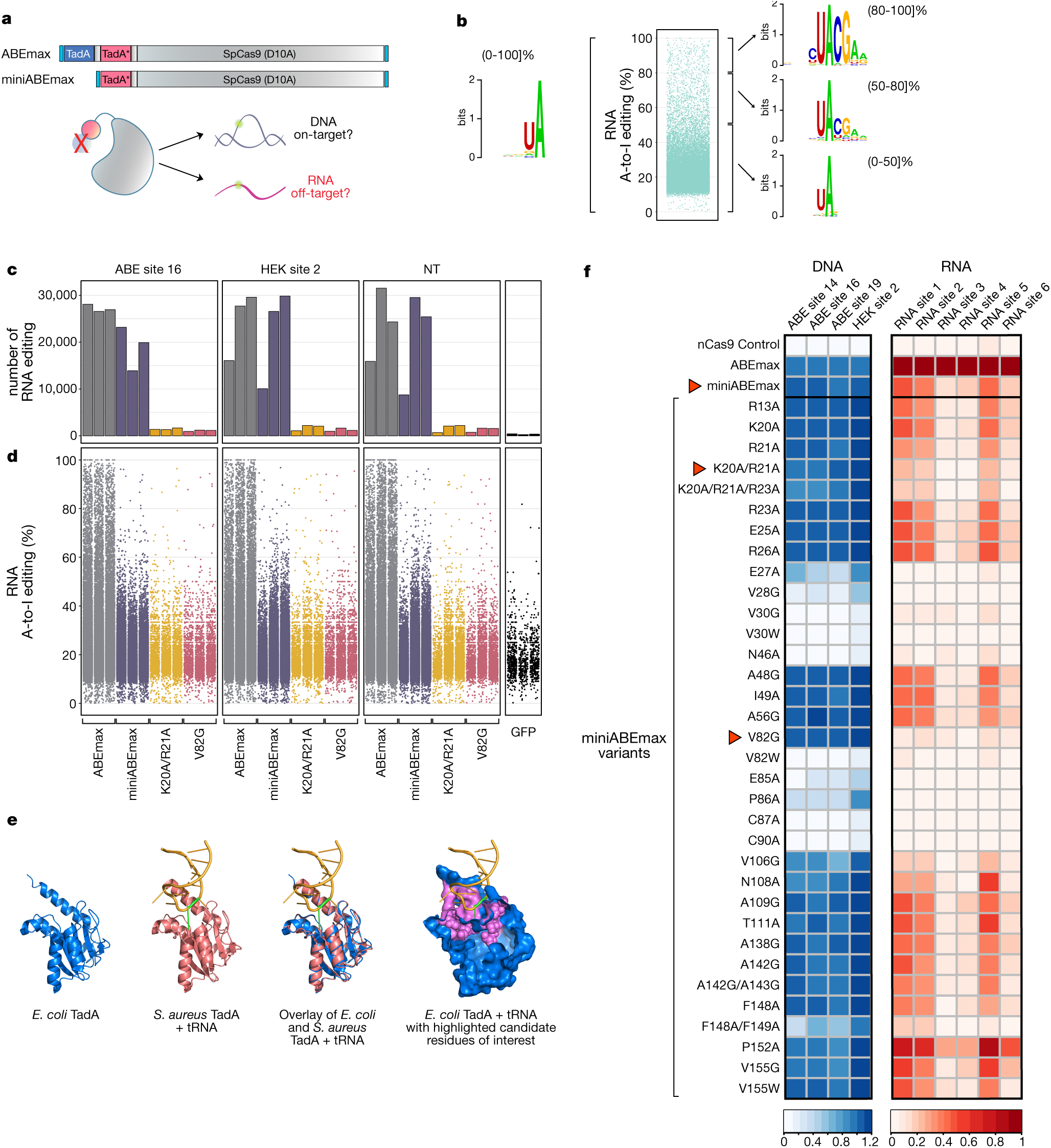
Engineering of SECURE-ABE variants with reduced off-target RNA editing activities. (a) Schematic illustration of ABEmax and miniABEmax architectures and overview of experimental testing of miniABEmax for on-target DNA and off-target RNA editing. Light blue boxes = bipartite NLS at N-and C-termini, TadA* = mutant TadA 7.10^3^, and small grey boxes = 32AA flanked XTEN linkers. nCas9 (SpCas9 D10A) = grey shape, TadA WT and mutant monomers = blue and red circles. Green halo = sites of potential adenine deamination on DNA and RNA. (b) Unstratified sequence logo (left) and stratified sequence logos for RNA adenines edited with high (80-100]%, middle (50-80]%, and low (0-50]% efficiencies by ABEmax. RNA-seq data shown in the Jitter plot was obtained from HEK293T cells in an earlier published study^5^. (c) Bar plots showing the number of RNA A-to-I edits observed in RNA-seq experiments in HEK293T cells with expression of ABEmax, miniABEmax, miniABEmax-K20A/R21A, or miniABEmax-V82G each with three different gRNAs (ABE site 16, HEK site 2, and non-targeting (NT)) and performed in independent biological replicates (n = 3). GFP negative control also performed in independent biological replicates (n = 3) is also shown. (d) Jitter plot showing the efficiencies of RNA A-to-I edits from the RNA-seq experiments shown in c. Each dot represents an edited adenine position in RNA. (e) Structural representations of *E. coli* TadA (PDB 1Z3A), structural representation of *S. aureus* TadA in complex with tRNA (PDB 2B3J), overlaid structures from *E. coli* TadA and *S. aureus* TadA, and surface representation of *E. coli* TadA in blue with backbone carbons of amino acid positions proximal to the predicted deaminase catalytic site highlighted in pink. Target adenine on tRNA (A34) marked in green. All graphical representations generated with PyMol (**Methods**). (f) Testing of 34 miniABEmax variants for their on-target DNA editing (A-to-G) and off-target RNA editing (A-to-I) activities. On-target DNA editing was assessed with four different gRNAs and off-target RNA alterations were screened on six RNA adenines previously identified as being efficiently modified by ABEmax^5^. Efficiencies are shown in heat map format (log-fold changes), with each box representing the mean of four independent biological replicates normalized to the edit efficiency observed with ABEmax for each target DNA or RNA site. Red arrows indicate three variants that were chosen for further analysis. Amino acid abbreviations are according to IUPAC nomenclature and residue numbering is based on the amino acid position in *E. coli* TadA. A = adenine; I = inosine. ABEmax = codon optimized adenine base editor. miniABEmax = ABEmax without N-terminal wild type TadA domain and the proximal 32AA linker.

We used RNA-seq to compare the transcriptome-wide off-target RNA editing activities of miniABEmax to ABEmax in HEK293T cells. Both editors and a nickase Cas9 (nCas9) control were each assayed in biological triplicate with three different gRNAs: two targeted to endogenous human gene sites (HEK site 2 and ABE site 16)^3^ and one to a site that does not occur in the human genome (NT)^5^. We performed these studies by sorting for GFP-positive cells (ABEmax was expressed as a P2A fusion with the base editor or nCas9 (**Methods**)). As an internal control, we first confirmed that ABEmax and miniABEmax both induced comparable on-target DNA editing efficiencies with HEK site 2 and ABE site 16 gRNAs (**Extended Data Fig. 1**). Edited RNA adenines were identified as previously described^5^ by filtering out background editing observed with read-count-matched negative controls (**Methods**). Surprisingly, the total number of edited adenines induced with miniABEmax expression was not consistently lower than what we observed with ABEmax --the two editors induced on average 80-fold and 54-fold more edited adenines relative to background (determined with a GFP-only negative control) (**Fig. 1c** and **Extended Data Table 1**). However, the overall distribution of individual RNA adenine edit efficiencies induced by miniABEmax were generally shifted to lower values (**Fig. 1d** and **Extended Data Fig. 2**). In addition, the sequence logos of the adenines edited by miniABEmax now appear to be shorter GUA or UA motifs, in contrast to the more extended CUACGAA motif characteristic of ABEmax (**Extended Data Fig. 3a** and **3b**).

We reasoned we might further reduce the off-target RNA editing activity of miniABEmax by altering amino acid residues within the remaining engineered *E. coli* TadA domain that could potentially mediate RNA recognition. However, although a crystal structure of isolated *E. coli* TadA has previously been solved^8^ (PDB 1Z3A; **Fig. 1e**), no structural information was available to delineate how this protein might recognize its RNA substrate. To overcome this, we exploited the availability of a *S. aureus* TadA-tRNA co-crystal structure^9^ (PDB 2B3J) (**Fig. 1e** and **Methods**). Although *E. coli* and *S. aureus* TadA share only partial amino acid sequence homology (39.5% identity; **Extended Data Fig. 4a**), we found that these two proteins share a high degree of structural homology (**Fig. 1e**). This similarity enabled us to overlay the two structures and thereby to infer 31 amino acid residues in *E. coli* TadA that likely lie near the enzymatic pocket around the substrate tRNA (**Fig. 1e**). In addition, we mutated three positively charged residues (R13, K20, and R21) in TadA* that we imagined might make contacts to the phosphate backbone of an RNA substrate.

We generated a total of 34 miniABEmax variants bearing various substitutions at the amino acid positions described above and screened each editor for on-target DNA editing and off-target RNA editing activities in HEK293T cells. To assess on-target DNA editing, we examined the efficiencies of A-to-G edits induced by each of the 34 variants with four gRNAs targeted to different endogenous gene sequences. To screen for off-target RNA editing activities, we quantified editing by each of the 34 variants at six RNA adenines using standard plasmid expression conditions (i.e., without sorting for GFP expression; see **Methods**); these six adenines were previously identified as being highly edited with ABEmax overexpression in HEK293T cells^5^. These experiments revealed that 23 of the 34 variants induced robust on-target DNA editing at least comparable to that observed with miniABEmax and ABEmax (**Fig. 1f** and **Extended Data Fig. 4b**). In addition, 14 of the 34 variants showed reduced editing activities on at least three of the six RNA adenines we examined relative to that observed with miniABEmax (**Fig. 1f, Extended Data Fig. 4b**). Importantly, three of the 34 variants (miniABEmax-K20A/R21A, -K20A/R21A/R23A, and -V82G) showed both robust on-target DNA editing activity and substantially reduced RNA editing activities (**Fig. 1f** and **Extended Data Fig. 4b and Extended Data Table 2**). Based on their DNA/RNA editing ratios, we chose to carry forward two miniABEmax variants (K20A/R21A and V82G) for more comprehensive characterization.

We characterized the transcriptome-wide off-target RNA editing profiles of the miniABEmax K20A/R21A and V82G variants using RNA-seq. The two variants were assessed in biological triplicate with the HEK site 2, ABE site 16, and NT gRNAs. In contrast to what we observed with miniABEmax, the K20A/R21A and V82G variants both induced substantially reduced numbers of edited adenines relative to ABEmax but still approximately four-fold and three-fold higher numbers, respectively, than background (determined with the GFP-only negative control) (**Fig. 1c**). In addition, the distribution of individual RNA adenine editing efficiencies for the two variants was shifted lower with both variants relative to ABEmax and miniABEmax (**Fig. 1d** and **Extended Data Fig. 2**). Overall, these results demonstrate, as we previously showed with the BE3 CBE, the feasibility of separating unwanted off-target RNA editing from desired on-target DNA editing activities with an ABE.

Having previously shown that off-target RNA editing occurs with BE3 CBE harboring the APOBEC1 enzyme^5^, we wanted to determine whether CBEs harboring other cytosine deaminases such as hA3A^10^, eA3A^11^ (an engineered A3A with more precise and specific DNA editing activities), or hAID^12^ might also induce unwanted edits. To do this, we transfected HEK293T cells in biological triplicates with plasmids expressing each of these CBEs and a guide RNA (gRNA) targeting a site in the *RNF2* gene. We then sorted cells with high CBE expression (top 5% of GFP signal) for isolation of genomic DNA (for on-target DNA amplicon sequencing) and total RNA (for RNA-seq) (**Methods**). Consistent with previously published studies^10-12^, hA3A-BE3 and eA3A-BE3 both showed robust on-target DNA editing (means of 91% and 82%, respectively, on position C6) with eA3A-BE3 showing greater precision while hAID-BE3 showed somewhat less efficient on-target DNA editing (mean of 32% on C6) (**Fig. 2a**). RNA-seq experiments revealed that hA3A-BE3 induced tens of thousands of C-to-U edits (**Fig. 2b** and **Extended Data Table 1**) distributed throughout the transcriptome (**Extended Data Fig. 5**). A number of these cytosines were edited with very high (>80%) efficiencies (**Fig. 2c** and **Extended Data Fig. 5**). Sequence logos derived from all cytosines edited by hA3A-BE3 show a consensus UC motif (**Extended Data Fig. 5**). However, sequence logos from subsets of Cs stratified by editing efficiencies reveal a more extended consensus sequence of CCAU<u>C</u>R for those Cs edited at higher efficiencies (**Extended Data Fig. 5**). This extended motif is consistent with a previous study that characterized RNA cytosines edited by the hA3A enzyme^13^. By contrast, eA3A-BE3 showed a dramatically reduced number of RNA edits relative to hA3A-BE3 but still slightly more (average of approximately three-fold) than what was observed with background in the GFP-only negative control (**Fig. 2b, Extended Data Fig. 6** and **Extended Data Table 1**). Interestingly, AID-BE3 showed numbers of RNA C-to-U edits comparable to what was observed in the negative control (**Fig. 2b, Extended Data Fig. 6** and **Extended Data Table 1**), consistent with a previous study that showed overexpression of isolated AID enzyme in activated B cells did not yield evidence for RNA editing^14^.

**Figure 2.**
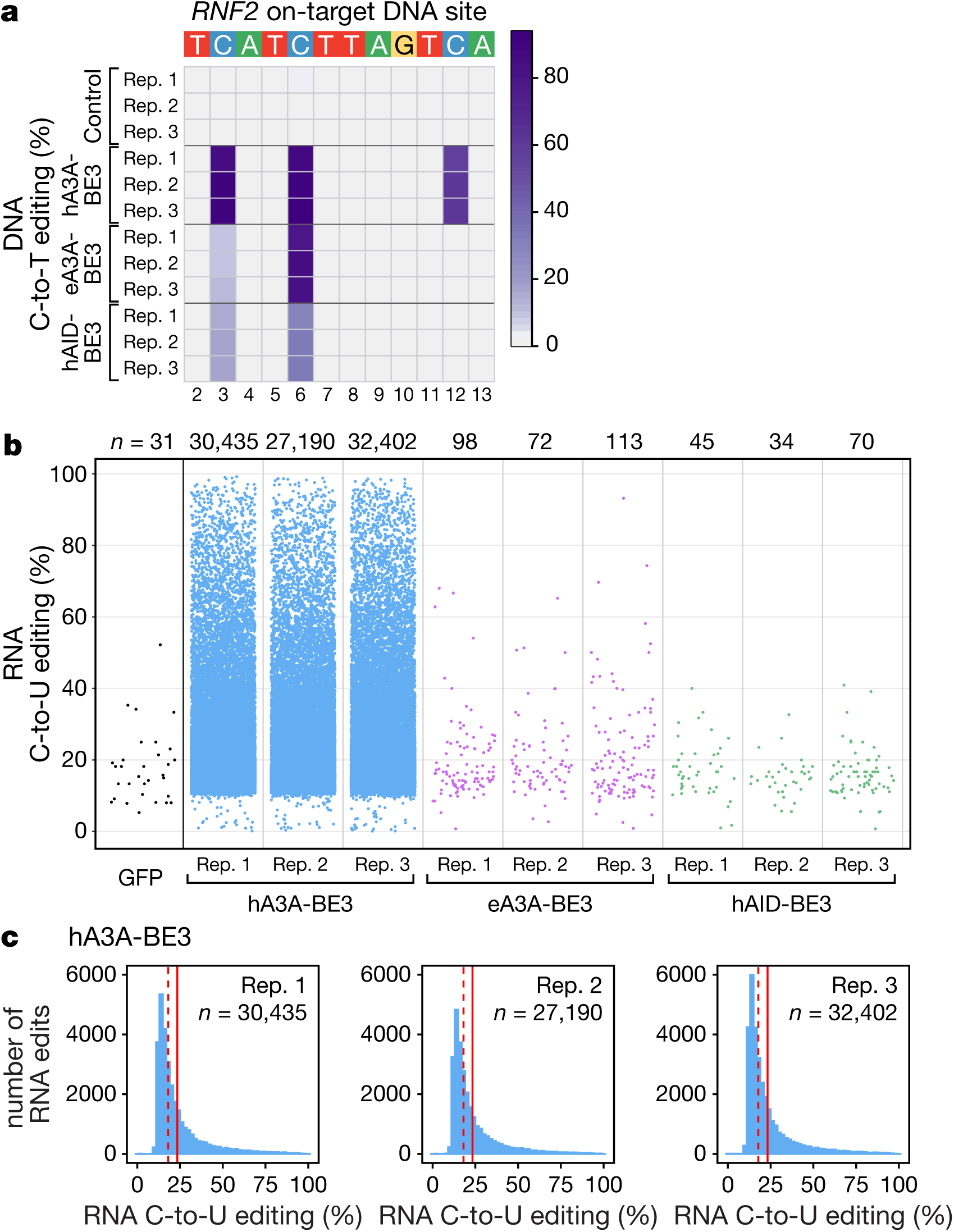
Transcriptome-wide off-target RNA editing activities of CBEs with non-APOBEC1 cytidine deaminases in HEK293T cells. (a) Heat map showing the on-target DNA editing efficiencies of hA3A-BE3, eA3A-BE3, hAID-BE3, and nCas9-UGI (Control) with a gRNA targeted to the *RNF2* gene (n= 3 independent replicates). Editing window shown includes only the most highly edited cytosines and not the entire spacer sequence. Numbering at the bottom represents spacer position with 1 being the most PAM distal location. (b) Jitter plots showing transcriptome-wide RNA C-to-U edits observed with a GFP negative control (single replicate) and hA3A-BE3, eA3A-BE3, and hAID-BE3 (each n= 3 biological replicates). Each dot represents a single edited cytosine. All experiments (except for the GFP control) were performed with co-expression of the *RNF2*-targeted gRNA and in all experiments the cells were sorted for top 5% of GFP/FITC signal except for the GFP control which was sorted for equivalent MFI of top 5% BE3 (**Methods**)). n = total number of modified cytosines. (c) Histograms showing the total number of RNA C-to-U edits observed (y-axis) for different editing efficiencies (x-axis) for the hA3A-BE3 RNA-seq experiments shown in b. n = total number of modified cytosines.

## Discussion

The work described here extends our understanding of the off-target RNA editing activities of DNA base editors, expands the options available to minimize these unwanted effects, and provides novel SECURE base editor architectures with desirable properties. The successful engineering of SECURE-ABE variants shows that, as we previously found with the BE3 CBE^5^, it is possible to minimize unwanted RNA editing while retaining efficient on-target DNA editing for an ABE. In the process of engineering these variants, we discovered a more extended consensus sequence motif for adenines edited with high efficiencies by ABEmax (CU<u>A</u>CGAA) that appears to be recognized by the wild-type TadA part of this fusion. Deletion of this TadA domain abolished recognition of these high efficiency sites and also resulted in the generation of the smallest SpCas9 base editors (1605 amino acids in length) described to date. In addition, our characterization of additional CBEs with deaminases other than APOBEC1 further expands the toolbox of base editors that can be used without inducing high-level RNA editing.

## Extended Data Figure and Table Legends

**Extended Data Fig. 1. On-target DNA editing activities of ABEmax, miniABEmax, and SECURE-ABE variants in HEK293T cells**

Heat maps showing the on-target DNA editing efficiencies of nCas9 (Control), ABEmax, miniABEmax, miniABEmax-K20A/R21A, and miniABEmax-V82G each assessed with two gRNAs targeted to ABE site 16 and HEK site 2 and performed in triplicate. Note that these were performed with the same transfected cells used for the RNA-seq experiments shown in **Figs. 1c and d**). Editing windows shown include only the most highly edited adenines and not the entire spacer sequence. Numbering at the bottom represents spacer position with 1 being the most PAM distal location.

**Extended Data Fig. 2. Additional data on the off-target RNA editing activities of ABEmax, miniABEmax, and SECURE-ABE in HEK293T cells**

Histograms showing the total number of RNA A-to-I edits observed (y-axis) for different editing efficiencies (x-axis) for ABEmax, miniABEmax, miniABEmax-K20A/R21A, and miniABEmax-V82G each tested with the ABE site 16, HEK site 2, and NT gRNAs. n= number of modified adenines. Experiments were performed in biological triplicate (data is derived from the same experiments as **Fig. 1c and d**). Dashed line = median; solid line = mean. Rep. = Replicate.

**Extended Data Figure 3. Sequence logos for RNA adenines edited by ABEmax and miniABEmax in HEK293T cells**

Sequence logos derived using all RNA-edited adenines (0-100]% or stratified RNA-edited adenines with high (80-100]%, middle (50-80]%, or low (0-50]% edit efficiencies induced by (a) ABEmax co-expressed with an ABE site 16, HEK site 2 or NT (non-targeting) gRNA or (b) miniABEmax co-expressed with an ABE site 16, HEK site 2, or NT gRNA. Logos are shown for biological triplicates from the same RNA-seq experiments displayed **Fig. 1c and d**. n = total number of modified adenines. For strata that contained <25 edited adenines, we considered the motif analysis as not sufficiently powered and therefore presented these logos in a semi-transparent fashion.

**Extended Data Figure 4. Engineering of miniABEmax variants with reduced off-target RNA adenine editing activities and preserved on-target DNA editing activities**

(a) Alignment of *E. coli* and *S. aureus* TadA amino acid sequences showing 39.5% identity (Uniprot alignment function). Stars represent 66 identical residues, dots and colons represent 54 residues of lower and higher similarity. Negative charges highlighted in red, positive charges highlighted in green. (b) Bar graphs of the data shown in **Fig. 1f**. On-target DNA editing (blue bar graphs) and off-target RNA editing (red bar graphs) efficiencies observed with negative control, ABEmax, miniABEmax, and 34 miniABEmax variants are shown for four on-target DNA sites (left panel, linear y-axis scale) and six RNA adenines previously shown to be edited by ABEmax (middle panel with linear y-axis scale and right panel with log10 y-axis scale). Means of four replicates are shown with individual quadruplicate biological replicate values (n = 4) overlaid as dots and error bars representing S.E.M. A = adenine; G = guanine; I = inosine.

**Extended Data Figure 5. Additional data showing transcriptome-wide off-target RNA editing activities and sequence logos of hA3A-BE3 in human HEK293T cells**

Manhattan plots showing transcriptome-wide distribution of RNA edits induced by hA3A-BE3 (these are the same data shown as Jitter plots in **Fig. 2b**). Sequence logos derived using all RNA-edited cytosines (0-100]% or stratified RNA-edited cytosines with high (80-100]%, middle (50-80]%, or low (0-50]% edit efficiencies induced by hA3A-BE3 expressed with the *RNF2* gRNA. n = total number of modified cytosines.

**Extended Data Figure 6. Histograms showing off-target RNA editing activity of eA3A-BE3 and hAID-BE3 in human HEK293T cells**

Histograms showing the total number of RNA C-to-U edits observed (y-axis) at different editing efficiencies (x-axis) with expression of eA3A-BE3 or hAID-BE3 co-expressed with the *RNF2* gRNA. Data from triplicate biological experiments are shown (derived from the data shown as Jitter plots in **Fig. 2b**). Dashed line = median; solid line = mean. Rep. = Replicate. n = number of modified cytosines.

**Extended Data Table 1. Summary of numbers of RNA edits observed in all RNA-seq experiments**

**Extended Data Table 2. Statistical tests for data on miniABEmax variant activities**

## Methods

### PyMOL Analysis of TadA structures

*Escherichia coli* tRNA-specific adenosine deaminase (TadA, PDB 1Z3A) and *Staphylococcus aureus* TadA with tRNA (PDB 2B3J) structures were downloaded from the Protein Data Bank and visualized with PyMOL version 2.2.2. Subunit A (monomer) of *S. aureus* TadA with tRNA was superimposed with subunit A of *E. coli* TadA using the “super” command. All figures were generated with PyMOL (<u>Schrödinger</u>).

### Plasmid cloning

All ABE constructs (reported in **Supplementary Table 1**) were cloned using the backbone and the P2A-EGFP-NLS fragment of ABEmax-P2A-EGFP-NLS (AgeI/NotI digest; Addgene ID 112101). ABEmax and variants were expressed under the control of a pCMV promoter. All CBE constructs were cloned using the backbone of SQT817 and expressed under the control of a pCAG promoter (AgeI-NotI-EcoRV digest, Addgene ID 53373). For the P2A-EGFP fragments in these constructs, we used BPK4335 (pCMV-BE3-P2A-EGFP) as a template. Guide RNA (gRNA) plasmids were cloned using the SpCas9 gRNA entry vector BPK1520 (pUC19 backbone; BsmbI cassette, Addgene ID 65777). All remaining constructs were generated using isothermal amplification (Gibson assembly, NEB). All gRNA and ABE plasmids were midi or maxi prepped using the Qiagen Midi/Maxi Plus kits.

### Cell culture

HEK293T cells (CRL-3216) and HepG2 cells (HB-8065) were purchased from and STR-authenticated by ATCC. Cells were cultured in Dulbecco’s Modified Eagle Medium (DMEM, Gibco) supplemented with 10% (v/v) fetal bovine serum (FBS, Gibco) and 1% (v/v) penicillin-streptomycin (Gibco) or Eagle’s Minimum Essential Medium with 10% (v/v) FBS and 0.5% (v/v) penicillin. Cells were passaged every 2-3 days when reaching around 80-90% confluency. Both cell lines were used only until passage 20 for all experiments, and the media was tested every two weeks for mycoplasma.

### Transfections

For ABE DNA on-target screening experiments, 2×10^4^ HEK293T cells were seeded into 96-well Flat Bottom Cell Culture plates (Corning), transfected 24h post seeding with 165ng base editor or negative control (bpNLS-32AA linker-nCas9(D10A)-bpNLS), 55ng guide RNA expression plasmid, and 0.66µL TransIT-293 (Mirus), and harvested 72h after transfection for DNA. For ABE RNA off-target screening experiments, 2×10^5^ HEK293T cells were seeded into 12-well Cell Culture plates (Corning), transfected 24h post seeding with 1.65µg base editor or negative control, 0.55µg guide RNA, and 6.6µL TransIT-293, and harvested 36h after transfection for RNA. For experiments with FACS-sorted cells, 6.5-7×10^6^ HEK293T cells were seeded into 150mm Cell Culture dishes (Corning), transfected 24h post seeding with 37.5µg base editor or an appropriate negative control fused to P2A-EGFP, 12.5µg guide RNA, and 150µL TransIT-293. Sorting took place 36-40h post transfection.

### Fluorescence-activated cell sorting (FACS)

Cells were prepared for sorting by diluting to 1×10^7^ cells per ml with 1X Phosphate Buffer Saline (PBS, Corning) supplemented with 10% FBS and filtering through 35µm cell strainer caps (Corning). Cells were sorted on a FACSAria II (BD Biosciences) using FACSDiva version 6.1.3 (BD Biosciences) after gating for single live cells. Cells treated with base editor were sorted for either all GFP signal (standard expression) or top 5% of cells with the highest GFP (FITC) signal (overexpression) into FBS; cells treated with nCas9 negative controls were sorted for either all GFP positive cells or the 5% of cells with a mean fluorescence intensity (MFI) matching that of the top 5% of cells treated with base editor. The GFP control shown in **Fig. 2b** was sorted to match the top 5% GFP signal of BE3-transfected control cells from the same day.

### DNA extraction

For ABE DNA on-target experiments, cells were lysed for DNA 72h post-transfection with freshly prepared 43.5µL DNA lysis buffer (50mM Tris HCl pH 8.0, 100mM NaCl, 5mM EDTA, 0.05% SDS, adapted from ref. 15), 5.25µL Proteinase K (NEB), and 1.25µL 1M DTT (Sigma). For experiments with sorted cells, cells were centrifuged (200*g*, 8 min) and lysed with 174µL DNA lysis buffer, 21µL Proteinase K, and 5µL 1M DTT. Lysates were incubated at 55°C on a plate shaker overnight, then gDNA were extracted with 2× paramagnetic beads (as described in ref. 16), washed 3 times with 70% EtOH, and eluted in 30µL 0.1X EB buffer (Qiagen).

### RNA extraction & reverse transcription

Cells were lysed for RNA 36h-40h post-transfection with 350µL RNA lysis buffer LBP (Macherey-Nagel), and RNA were extracted with the NucleoSpin RNA Plus kit (Macherey-Nagel) following the manufacturer’s instructions. RNA was then reverse transcribed into cDNA with the High Capacity RNA-to-cDNA kit (Thermo Fisher) following the manufacturer’s instructions.

### Library preparation for DNA or cDNA targeted amplicon sequencing

Next-generation sequencing (NGS) of DNA or cDNA was performed as previously described^5^. In summary, the first PCR was performed to amplify genomic or transcriptomic sites of interested with primers containing Illumina forward and reverse adapter sequences (see **Supplementary Table 2** for primers and amplicons used in this study), following NEB Phusion High-Fidelity DNA Polymerase instructions. The first PCR products were cleaned with 0.7x paramagnetic beads, then the second PCR was performed to add barcodes with primers containing unique sets of p5/p7 Illumina barcodes (analogous to TruSeq CD indexes). The second PCR products were again cleaned with 0.7× paramagnetic beads. The libraries were then pooled based on concentrations measured with the QuantiFluor dsDNA System (Promega) and Synergy HT microplate reader (BioTek) at 485/528nm. The final pool was quantified by qPCR with the NEBNext Library Quant Kit for Illumina (NEB) and sequenced paired-end (PE) 2×150 on the Illumina MiSeq machine using 300-cycle MiSeq Reagent Kit v2 or Micro Kit v2 (Illumina). FASTQs (post-demultiplexing) were downloaded from Illumina BaseSpace and analyzed using a batch version of CRISPResso 2.

### RNA library preparation & sequencing

RNA-seq experiments were performed as previously described^5^. Briefly, RNA libraries were prepared with the TruSeq Stranded Total RNA Library Prep Gold kit (Illumina) following the manufacturer’s instructions. SuperScript III (Invitrogen) was used for first-strand synthesis, and IDT for Illumina TruSeq RNA unique dual indexes (96 indexes) were used to avoid index hopping. The libraries were pooled based on qPCR measurements with the NEBNext Library Quant Kit for Illumina. The final pool was sequenced PE 2×76 on the Illumina HiSeq2500 machine (for all CBE experiments and one ABE experiment shown in Fig. 1b) or PE 2×100 on the NovaSeq6000 machine (for all remaining ABE experiments) at the Broad Institute of Harvard and MIT (Cambridge, MA). To account for variable sequencing depths, all RNA-seq libraries sequenced on the NovaSeq were uniformly downsampled to 100 million reads per library using seqtk version 1.0-r82-dirty (https://github.com/lh3/seqtk).

### Amplicon sequencing analysis

Amplicon sequencing data was analyzed with CRISPResso2 v.2.0.27^17^. The heatmaps for the SECURE-ABE screening in **Fig. 1f** display the highest edited adenine at the target site. Editing efficiency values were log2 transformed with a pseudocount of 1, averaged over quadruplicates, and normalized to ABEmax. The remaining heatmaps showing ABE and CBE on-target DNA editing (**Fig. 2** and **Extended Data Fig. 1**) show an editing window that includes the edited Cs and a grey background for editing efficiencies smaller than 2%. Tables with the full output will be made available as Supplementary Materials.

### RNA variant calling pipeline

All bioinformatic analysis was performed in concordance with GATK Best Practices^18,19^ for RNA-seq mutation calling as we have previously described^5^. Briefly, raw sequencing reads were two-pass aligned to the reference hg38 reference genome with STAR^20^ with parameters to discard multi-mapping reads. After PCR duplicate removal and base recalibration, mutations in RNA-seq libraries were called using GATK HaplotypeCaller. RNA edits in CBE and ABE overexpression experiments were identified using a downstream modification of the GATK pipeline output as we have previously described^5^. Specifically, mutation positions called by HaplotypeCaller were further filtered to include only those satisfying the following criteria with reference to the corresponding control experiments: (1) Read coverage for a given edit in control experiment should be greater than the 90th percentile of read coverage across all edits in the overexpression experiment. (2) 99% of reads covering each edit in the control experiment were required to contain the reference allele. Edits were further filtered to exclude those with fewer than 10 reads or 0% alternate allele frequencies. A-G edits include A-G edits identified on the positive strand as well as T-C edits identified on the negative strand. For CBE overexpression experiments, C-T edits include C-T edits identified on the positive strand as well as G-A edits derived from the negative strand.

Six A-to-I edits identified from the above pipeline were chosen to test SECURE ABE variants based on the following criteria. These were sites that had (1) read coverage of at least 50 in all replicates of control and overexpression experiments, (2) 99% reads in all control experiments containing reference allele and (3) at least 60% alternate allele frequencies in all replicates. From this list, primers were tested for the top 15 edited sites that were also within 150 bases of an exon-exon junction and the 6 highest edited sites with robust amplification from cDNA were chosen.

## Acknowledgements

J.K.J., J.G., and R.Z. are supported by the Defense Advanced Research Projects Agency (HR0011-17-2-0042). Support was also provided by the National Institutes of Health (RM1 HG009490 to J.K.J. and J.G. and R35 GM118158 to J.K.J. and M.J.A.). J.K.J. is additionally supported by the Desmond and Ann Heathwood MGH Research Scholar Award. We thank G. Ciaramella of Beam Therapeutics for the suggestion to delete the wild-type TadA monomer from ABEmax. We thank A. Lapinate of the Doudna Lab for suggesting the overlay of *E. coli* and *S. aureus* TadA structures and S.J. Lee for technical assistance.

