## Extended Data Figures and Tables for "CRISPR adenine and cytosine base editors with reduced RNA off-target activities"

#### Extended Data Figure 1

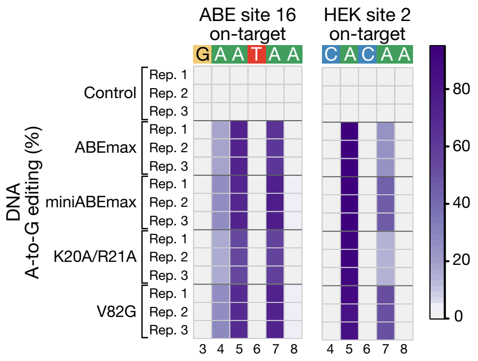

#### Extended Data Figure 2

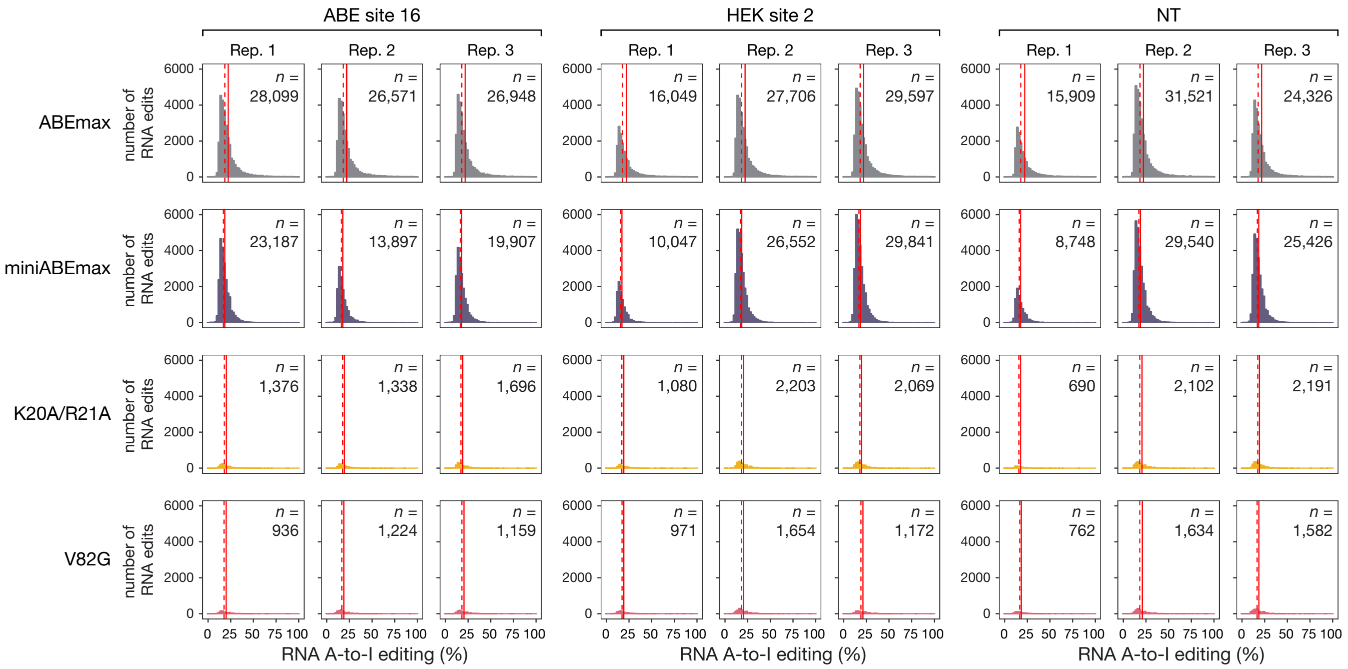

**Extended Data Figure 3**

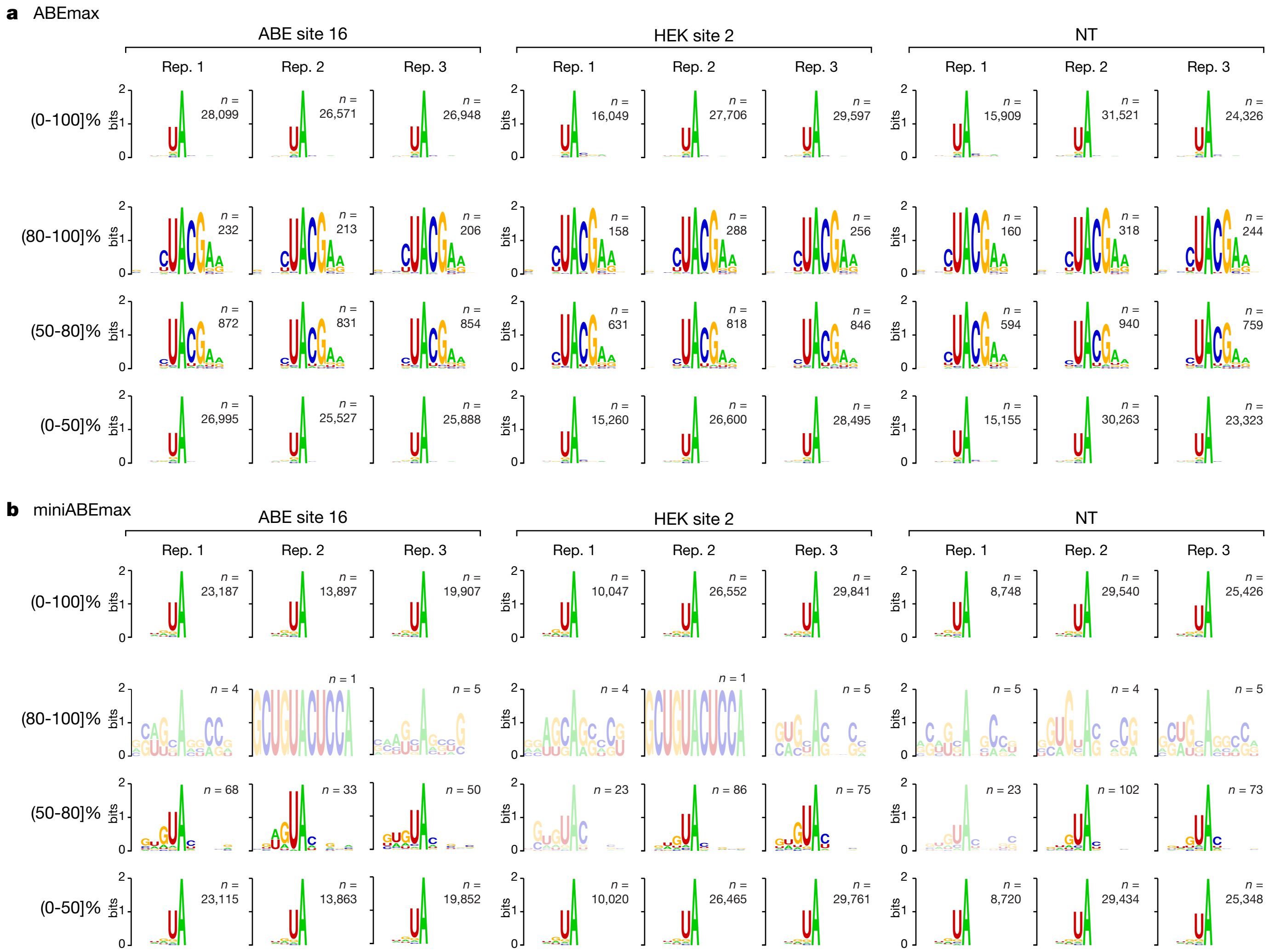

### Extended Data Figure 4

**a**

```
P68398 TADA_ECOLI     1    MSEVEFSHEYWMRHALTLAKRAWDEREVPVGAVLVHNNRVIGEGWNRPIGRHDPTAHAEI    60
Q99W51 TADA_STAAM     1    ----MTNDIYFMTLAIEEAKKAAQLGEVPIGAIITKDDEVIARAHNLRETLQQPTAHAEH    56
                                 .. *:*  *:  **:* :  ***:**::.:::.**... *     ::******

P68398 TADA_ECOLI    61    MALRQGGLVMQNYRLIDATLYVTLEPCVMCAGAMIHSRIGRVVFGARDAKTGAAGSLMDV   120
Q99W51 TADA_STAAM    57    IAIERAAKVLGSWRLEGCTLYVTLEPCVMCAGTIVMSRIPRVVYGADDPKGGCSGSLMNL   116
                           :*:.:.. *: .:** ..**************::: *** ***:** * * *.:****::

P68398 TADA_ECOLI   121    LHHPGMNHRVEITEGILADECAALLSDFFRMRRQEIKAQKKAQSSTD   167
Q99W51 TADA_STAAM   117    LQQSNFNHRAIVDKGVLKEACSTLLTTFFKNLRANKKSTN-------   156
                           *:: .:***. : :*:* : *::**: **:  * : *: :
```

**b**

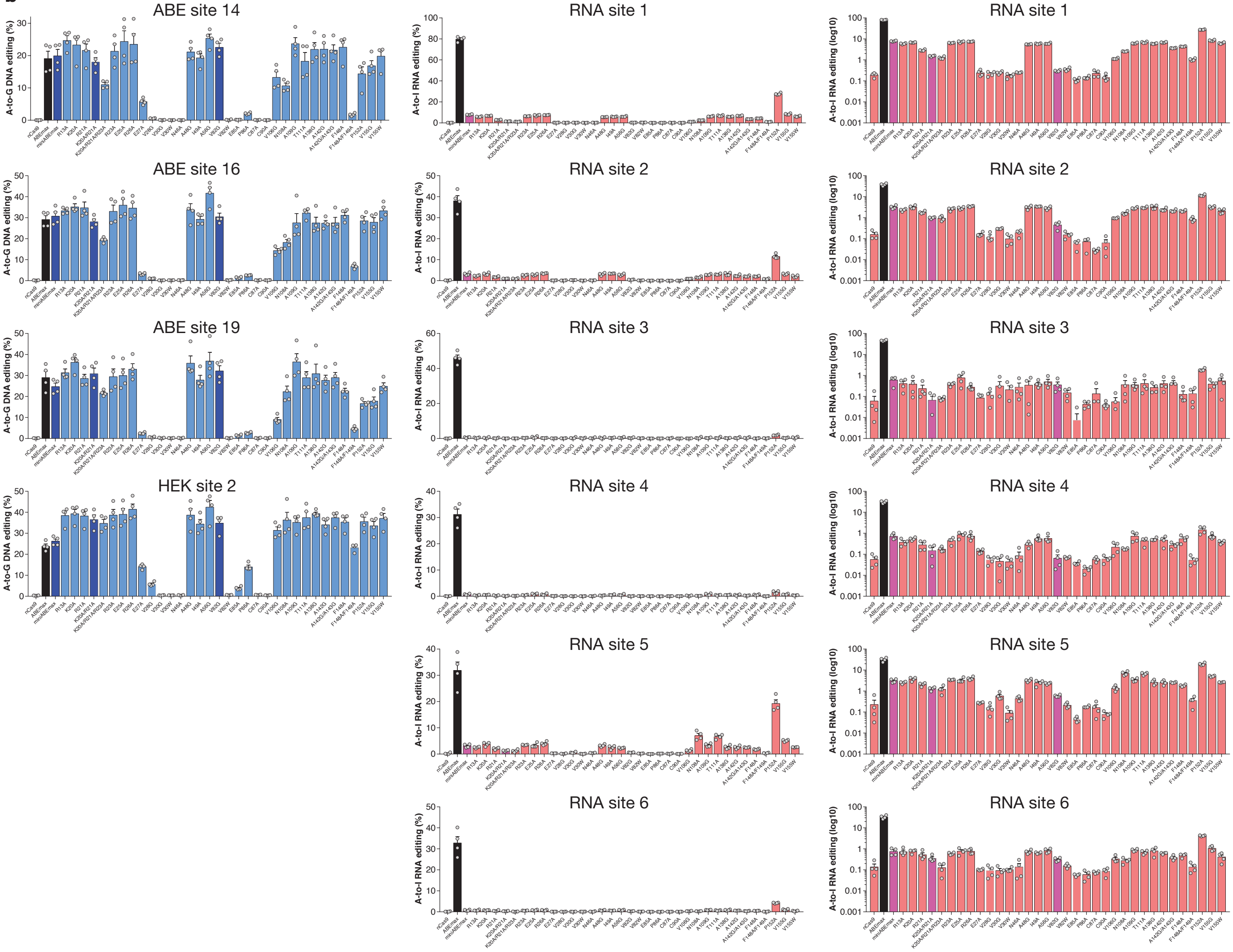

#### Extended Data Figure 5

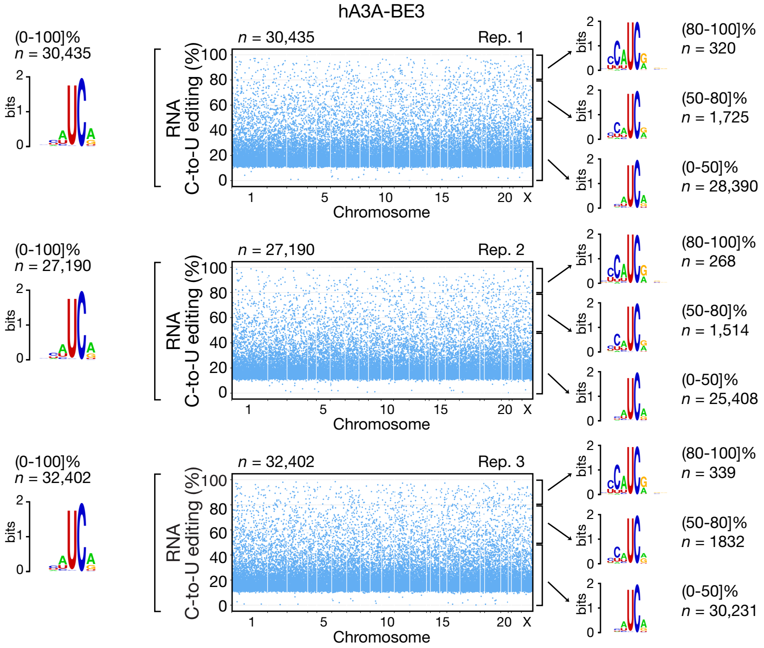

#### Extended Data Figure 6

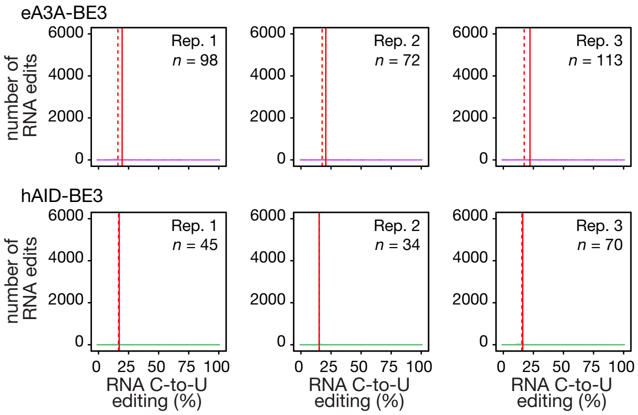

**Extended Data Table 1**

| Figure | Cell | BE | gRNA | Sort | Replicate | **A-to-I (for ABE) or C-to-U (for CBE)** | Other | A-to-I or C-to-U (%) |
|---|---|---|---|---|---|---|---|---|
| Fig. 1b | HEK293T | ABEmax | HEK site 2 | Top 5% | Rep. 1 | **37,061** | 88 | 99.763 |
| Fig. 1c & d | HEK293T | ABEmax | ABE site 16 | All GFP | Rep. 1 | **28,099** | 197 | 99.304 |
| | | | | | Rep. 2 | **26,571** | 238 | 99.112 |
| | | | | | Rep. 3 | **26,948** | 238 | 99.125 |
| | | miniABEmax | ABE site 16 | All GFP | Rep. 1 | **23,187** | 216 | 99.077 |
| | | | | | Rep. 2 | **13,897** | 202 | 98.567 |
| | | | | | Rep. 3 | **19,907** | 232 | 98.848 |
| | | miniABEmax-K20A/R21A | ABE site 16 | All GFP | Rep. 1 | **1,376** | 292 | 82.494 |
| | | | | | Rep. 2 | **1,338** | 291 | 82.136 |
| | | | | | Rep. 3 | **1,696** | 295 | 85.183 |
| | | miniABEmax-V82G | ABE site 16 | All GFP | Rep. 1 | **936** | 243 | 79.389 |
| | | | | | Rep. 2 | **1,224** | 336 | 78.462 |
| | | | | | Rep. 3 | **1,159** | 269 | 81.162 |
| | HEK293T | ABEmax | HEK site 2 | All GFP | Rep. 1 | **16,049** | 201 | 98.763 |
| | | | | | Rep. 2 | **27,706** | 246 | 99.120 |
| | | | | | Rep. 3 | **29,597** | 193 | 99.352 |
| | | miniABEmax | HEK site 2 | All GFP | Rep. 1 | **10,047** | 231 | 97.752 |
| | | | | | Rep. 2 | **26,552** | 251 | 99.064 |
| | | | | | Rep. 3 | **29,841** | 177 | 99.410 |
| | | miniABEmax-K20A/R21A | HEK site 2 | All GFP | Rep. 1 | **1,080** | 238 | 81.942 |
| | | | | | Rep. 2 | **2,203** | 383 | 85.189 |
| | | | | | Rep. 3 | **2,069** | 315 | 86.787 |
| | | miniABEmax-V82G | HEK site 2 | All GFP | Rep. 1 | **971** | 216 | 81.803 |
| | | | | | Rep. 2 | **1,654** | 333 | 83.241 |
| | | | | | Rep. 3 | **1,172** | 276 | 80.939 |
| | HEK293T | ABEmax | NT | All GFP | Rep. 1 | **15,909** | 202 | 98.746 |
| | | | | | Rep. 2 | **31,521** | 229 | 99.279 |
| | | | | | Rep. 3 | **24,326** | 196 | 99.201 |
| | | miniABEmax | NT | All GFP | Rep. 1 | **8,748** | 379 | 95.847 |
| | | | | | Rep. 2 | **29,540** | 244 | 99.181 |
| | | | | | Rep. 3 | **25,426** | 261 | 98.984 |
| | | miniABEmax-K20A/R21A | NT | All GFP | Rep. 1 | **690** | 206 | 77.009 |
| | | | | | Rep. 2 | **2,102** | 325 | 86.609 |
| | | | | | Rep. 3 | **2,191** | 265 | 89.210 |
| | | miniABEmax-V82G | NT | All GFP | Rep. 1 | **762** | 143 | 84.199 |
| | | | | | Rep. 2 | **1,634** | 304 | 84.314 |
| | | | | | Rep. 3 | **1,582** | 282 | 84.871 |
| | HEK293T | GFP | -- | All GFP | Rep. 1 | **423** | 202 | 67.680 |
| | | | | | Rep. 2 | **270** | 175 | 60.674 |
| | | | | | Rep. 3 | **363** | 168 | 68.362 |
| Fig. 2b | HEK293T | GFP | -- | MFI-matched to top 5% BE3 expression | Rep. 1 | **31** | 131 | 19.136 |
| | | hA3A-BE3 | *RNF2* | Top 5% | Rep. 1 | **30,435** | 8 | 99.974 |
| | | | | | Rep. 2 | **27,190** | 8 | 99.971 |
| | | | | | Rep. 3 | **32,402** | 11 | 99.966 |
| | | eA3A-BE3 | *RNF2* | Top 5% | Rep. 1 | **98** | 101 | 49.246 |
| | | | | | Rep. 2 | **72** | 87 | 45.283 |
| | | | | | Rep. 3 | **113** | 78 | 59.162 |
| | | hAID-BE3 | *RNF2* | Top 5% | Rep. 1 | **45** | 201 | 18.293 |
| | | | | | Rep. 2 | **34** | 144 | 19.101 |
| | | | | | Rep. 3 | **70** | 234 | 23.026 |

**Extended Data Table 2**

| DNA | ABEmax vs miniABEmax | ABEmax vs K20A/R21A | ABEmax vs V82G | miniABEmax vs K20A/R21A | miniABEmax vs V82G |
|---|---|---|---|---|---|
| ABE site14 | 0.78601 | 0.69378 | 0.23183 | 0.45035 | 0.29956 |
| ABE site16 | 0.58244 | 0.65370 | 0.62989 | 0.32954 | 0.90884 |
| ABE site19 | 0.27139 | 0.67921 | 0.45482 | 0.11499 | 0.05184 |
| HEK site2 | 0.16278 | 0.00461 | 0.00737 | 0.01031 | 0.01829 |

| RNA | ABEmax vs miniABEmax | nCas9 Control vs miniABEmax | ABEmax vs K20A/R21A | ABEmax vs V82G | miniABEmax vs K20A/R21A | miniABEmax vs V82G |
|---|---|---|---|---|---|---|
| RNA site1 | 0.00001 | 0.00003 | 0.00001 | 0.00001 | 0.00002 | 0.00004 |
| RNA site2 | 0.00067 | 0.00215 | 0.00064 | 0.00061 | 0.00523 | 0.00215 |
| RNA site3 | 0.00011 | 0.01602 | 0.00011 | 0.00011 | 0.01714 | 0.19767 |
| RNA site4 | 0.00063 | 0.00891 | 0.00061 | 0.00060 | 0.00865 | 0.00824 |
| RNA site5 | 0.00287 | 0.00115 | 0.00253 | 0.00239 | 0.00746 | 0.00419 |
| RNA site6 | 0.00182 | 0.01755 | 0.00178 | 0.00178 | 0.06036 | 0.05016 |

*p-values* generated with two-tailed t-test (type 3)
